# Gaze Bias Differences Capture Individual Choice Behavior

**DOI:** 10.1101/228825

**Authors:** Armin W. Thomas, Felix Molter, Ian Krajbich, Hauke R. Heekeren, Peter N. C. Mohr

**Author notes:** Shared first authorship with equal contribution (Armin W. Thomas, Felix Molter). Author Note: Armin W. Thomas, Electrical Engineering and Computer Science, Technische Universität Berlin; Felix Molter, School of Business & Economics, Freie Universität Berlin; Ian Krajbich, Department of Psychology, Department of Economics, Ohio State University; Hauke R. Heekeren, Department of Education & Psychology, Freie Universität Berlin; Peter N. C. Mohr, School of Business & Economics, Freie Universität Berlin. Ordering of shared first authorship was determined by the following coin toss procedure: A.W.T. brought three Canadian Dollar coins from which F.M. selected one. F.M. chose one of the coin’s sides. F.M. flipped the coin and let it fall onto a hard wooden floor. The upper side of the coin then determined the ordering outcome. Correspondence concerning this article should be addressed to Peter N. C. Mohr, School of Business & Economics, Freie Universität Berlin, Garystr. 21, 14195 Berlin, Germany.

## Abstract

How do we make simple consumer choices (e.g., deciding between an apple, an orange, and a banana)? Recent empirical evidence suggests a close link between choice behavior and eye movements at the group level, with generally higher choice probabilities for items that were looked at longer during the decision process. However, it is unclear how variable this effect is across individuals. Here, we investigate this question in a multialternative forced-choice experiment using a novel computational model that can be easily applied to the individual participant level. We show that a link between gaze and choice is present for most individuals, but differs considerably in strength, namely, the choices of some individuals are almost independent of gaze allocation, while the choices of others are strongly associated with gaze behavior. Accounting for this variability in our model allows us to explain and accurately predict individual differences in observed choice behavior.

---

In everyday life, we are constantly confronted with simple consumer choices, such as whether to have an apple or a banana for breakfast, or which bottle of juice to buy at the supermarket. Traditional models describing this type of consumer choice assume that people assign a utility (or subjective value) to each available option and make utility-maximizing choices (Von Neumann & Morgenstern, 1945). Notably, choices are assumed to be based solely on option attributes, and thereby, are independent of information search processes during the decision. This assumption has recently been challenged by a variety of empirical findings showing that the allocation of gaze during the decision-making process also plays a significant role, as a longer gaze towards one option is regularly associated with a higher choice probability for that option (Armel, Beaumel, & Rangel, 2008; Cavanagh, Wiecki, Kochar, & Frank, 2014; Fiedler & Glöckner, 2012; Folke, Jacobsen, Fleming, & De Martino, 2016; Glöckner & Herbold, 2011; Konovalov & Krajbich, 2016; Krajbich & Rangel, 2011; Krajbich, Armel, & Rangel, 2010; Krajbich, Lu, Camerer, & Rangel, 2012; Pärnamets et al., 2015; Shimojo, Simion, Shimojo, & Scheier, 2003; Stewart, Gächter, Noguchi, & Mullett, 2015; Stewart, Hermens, & Matthews, 2016; Vaidya & Fellows, 2015). Further, external manipulation of an individual’s gaze allocation changes choice probabilities accordingly (Armel et al., 2008; Pärnamets et al., 2015; Shimojo et al., 2003; Tavares, Perona, & Rangel, 2017).

These findings led to the development of novel computational models, which integrate eye movement data into the choice process and formalize the empirically observed association between gaze and choice (Ashby, Jekel, Dickert, & Glöckner, 2016; Cavanagh et al., 2014; Fisher, 2017; Krajbich & Rangel, 2011; Krajbich et al., 2010, 2012; Towal, Mormann, & Koch, 2013). These models are based on classic evidence accumulation models (Ratcliff, 1978; Ratcliff, Smith, Brown, & McKoon, 2016) and make the additional assumption that the momentary rate of evidence accumulation depends on the decision maker’s eye movements: Evidence accumulation for an option is assumed to be discounted by a constant factor while another item is fixated upon. Accounting for this gaze bias, these models provide a precise quantitative account of many aspects of simple consumer choice behavior at the group level.

While group level statistics are informative for some research questions (e.g., forecasting product sales in economic research), statements about the majority of people, or the “average person”, are often unsuitable for understanding the choice behavior of an individual. Even worse, aggregate models of behavior can lead to false conclusions about true underlying individual processes (Grandy, Lindenberger, & Werkle-Bergner, 2017; Lewandowsky & Farrell, 2010): In a learning task, for example, the group level average learning curve would appear as a gradual, smooth function over time, even if all individuals showed abrupt, step-like learning curves (much like an epiphany), but with variable learning onsets across individuals (Hayes, 1953). This group level model, however, would not describe any individual of the group well, and the deduction that individual learning occurs smoothly would be false. It is thus crucial to understand and explain choice behavior at the individual level.

Similarly, previously reported group level models quantifying the association between gaze and choice specified a constant gaze bias for all individuals (e.g., Krajbich et al., 2010; Krajbich & Rangel, 2011), without testing the model performance on the individual level. It therefore remains to be shown whether an association between gaze and choice is present across individuals and whether the strength of such an association is constant. If, however, people’s decisions were affected differently by looking behavior, we would find that the choices of some individuals are more biased by gaze, and therefore, more inconsistent with subjective value ratings than other individuals’ choices. Imagine, for example, a choice between two bottles of orange juice in the supermarket: one has a slightly higher utility for the decision maker than the other, but it is also less visually salient. If this person’s association of gaze and choice behavior was strong, her choice would be biased towards the bottle that is attracting more of her gaze, even though it has lower utility. On the other hand, if the person’s association is weak, she would then be able to select the option that is higher in utility, despite her gaze being attracted more towards the inferior option. Accordingly, if the strength of this association is variable across individuals, it is necessary to account for these differences to accurately predict individual choice behavior.

Here, we investigated whether the previously reported link between gaze and choice behavior is variable across individuals, using a novel computational model that can easily be applied to individual participant and multialternative choice data. With this model, we reaffirmed that an association between gaze and choice is present at the group level, and indeed, present for most individuals. The strength of this association, however, showed substantial variability. By accounting for this variability, we were able to explain and accurately predict empirically observed differences in individuals’ choice behavior.

## Results

### Data set & Task overview

To investigate individual differences in the influence of gaze allocation on simple economic choice behavior, we used a previously published, prototypical data set (Krajbich & Rangel, 2011) that we obtained from the original authors in a preprocessed format (see Methods for full details). In the corresponding experiment, hungry participants made choices between three snack food items, without time restrictions (Figure 1). Participants also gave a liking rating for each of the 70 snack food items that were used in the experiment. During the choice task, the participants’ eye movements were continuously recorded using an eye tracker. The data set included 30 participants, who performed 100 choice trials each.

**Figure 1:**
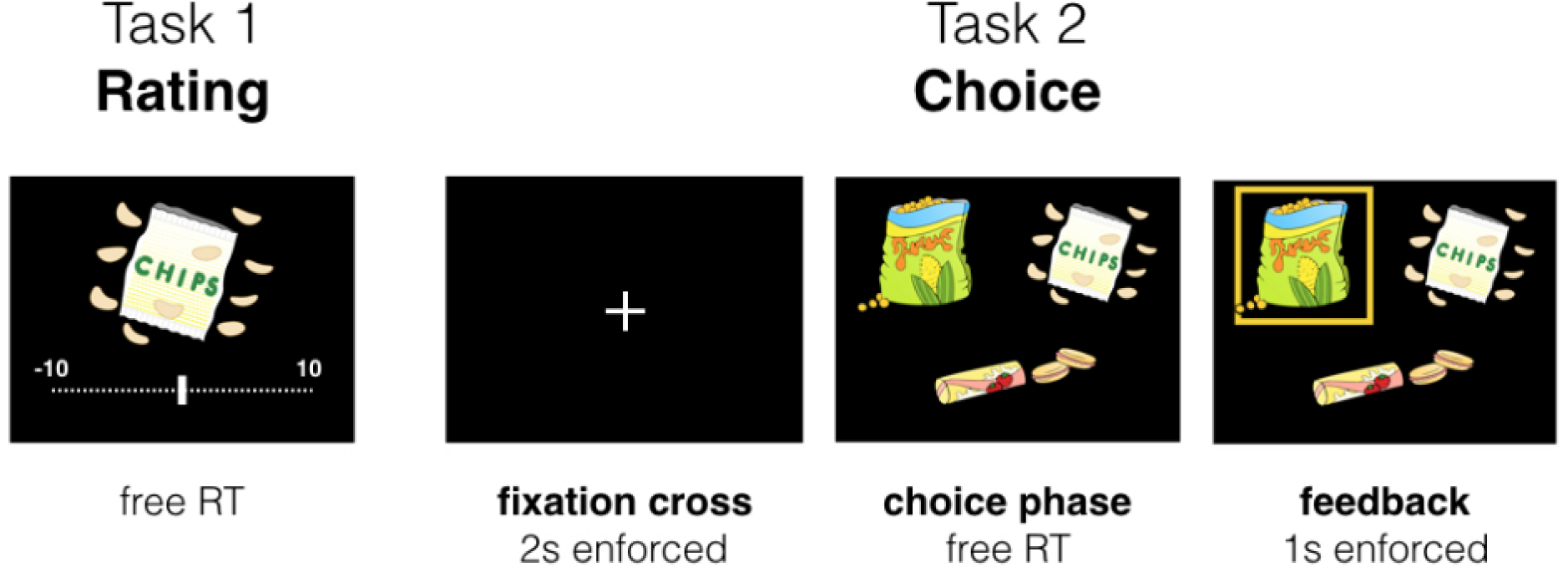
Experimental Paradigm. All the participants completed two tasks in a single session. **Task 1:** The participants rated all 70 snack foods items on a liking rating scale between −10 and 10, according to how much they would like to eat each item. **Task 2:** In each choice trial, the participants were required to maintain a central fixation for 2 s. Next, the participants were asked to choose the item that they would like to eat most from sets of 3 choice items, while their eye movements were being recorded. Choices were followed by 1 s of visual feedback.

### Individual differences in the data

We analyzed three metrics for individual differences, namely, (i) the participants’ response time, (ii) the mean probability of choosing the item with the highest liking rating, and (iii) the influence of gaze allocation on choice probability (mean increase in choice probability for an item that was fixated on longer than the others, after correcting for the influence of the item value). We found that participants differed considerably in all of the three metrics (Figure 2):

**Figure 2:**
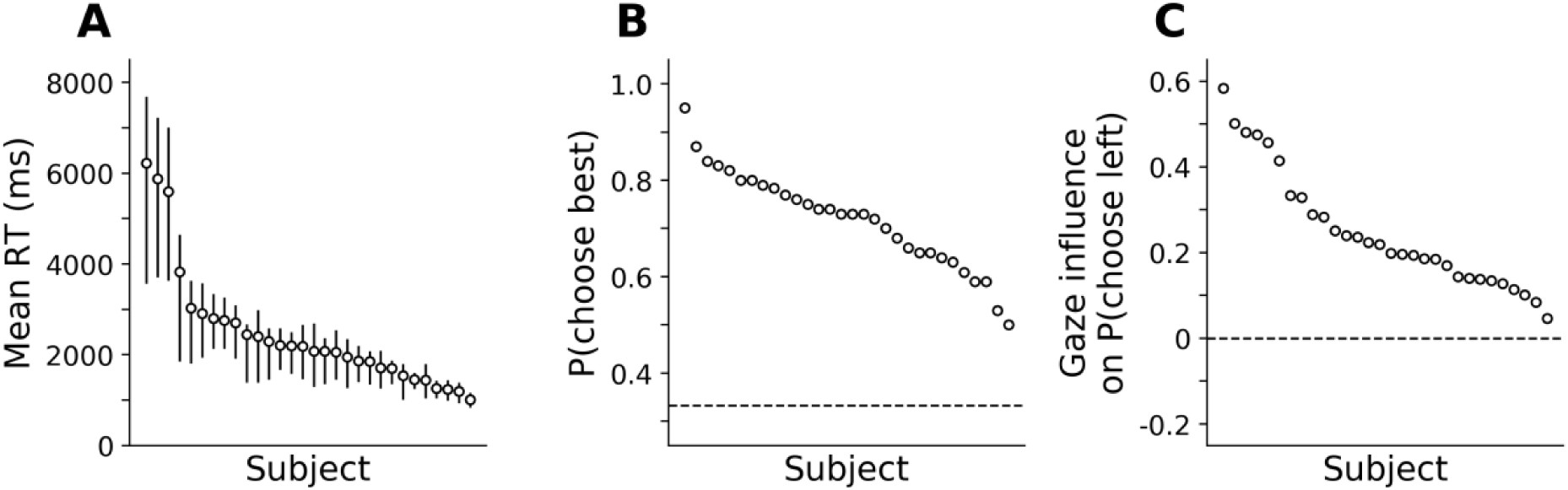
Individual differences in the three behavioral metrics: response time, probability of choosing the highest rated item and influence of gaze on choice probability. **A**: Mean individual response times (error bars denote first and third quartile). **B**: Mean probability of choosing the item with the highest liking rating. The dashed horizontal line indicates chance level accuracy. **C**: Individual influence of gaze on choice probability (mean increase in choice probability for an item that is fixated longer than the others, after correcting for the influence of item value). The data points are sorted from high to low in each panel.

The participants’ mean response times ranged from 1006 to 6217 ms, with mean ± s.d. = 2462 ± 1298 ms (Figure 2A), while their probabilities of choosing the highest rated item in a trial ranged from 50.00% to 95.00%, with mean ± s.d. = 71.94% ± 10.01% (Figure 2B).

We also probed the relationship between individual allocation of gaze and choice. Previous work in simple choice tasks has shown that individuals are, on average, more likely to choose an option when they spent more time fixating on it, relative to the others (Armel et al., 2008; Cavanagh et al., 2014; Folke et al., 2016; Krajbich & Rangel, 2011; Krajbich et al., 2010). Here, we devised an individual measure to quantify the relationship between gaze allocation and choice for each individual: following previous work (Krajbich & Rangel, 2011; Krajbich et al., 2010), for each participant we first estimated the probability of choosing the left item in a choice set using logistic regression, based on its relative item value (the difference between the item’s value and the mean value of all other items in that trial) and the range between the other items’ value. We then subtracted this estimated probability from the empirically observed choice (either 1 if the left item was chosen, or 0 otherwise). Finally, we averaged the resulting “residual” choice probability for trials in which the left item had a positive and negative final gaze advantage (computed as the difference in the fraction of the total fixation time that the participants spent fixating on the left item and the average fraction that they spent fixating on the others). The difference between these two described the average difference in choice probability for the items with a positive versus negative final gaze advantage, when corrected for the influence of relative item value on choice probability and the other items’ range of values. We found that individual scores on this measure ranged from 0.05 to 0.58, with mean ± s.d. = 0.25 ± 0.14 (Figure 2C). Notably, all the participants showed positive scores, indicating an overall positive relationship between gaze allocation and choice. We did, however, find strong variation in this measure.

### Modeling individual differences in simple economic choice

The behavioral and eye tracking data suggest substantial variability in the extent to which gaze affects a participant’s choice behavior (Figure 2C). The computational mechanism relating gaze and choice patterns, however, cannot be inferred from descriptive analyses alone. In addition, conclusive quantitative evidence for or against the presence of a mechanism that biases choices depending on the distribution of gaze has yet to be provided at the individual level. We therefore adopted a principled computational modeling approach to investigate whether a formalized gaze bias mechanism, in conjunction with individual gaze patterns, can improve model predictions of individual choice and response time data.

We propose a new model called the Gaze-weighted Linear Accumulator Model (GLAM; Figure 3) that is inspired by the attentional Drift Diffusion Model (aDDM) proposed by Krajbich et al. (2010; also, Krajbich & Rangel, 2011; Krajbich et al., 2012). The GLAM assumes accumulation of evidence in favor of each item, that is modulated by gaze behavior: While an item is not fixated on, accumulation occurs at a rate discounted by the gaze bias parameter γ. A choice is made as soon as evidence in favor of one item reaches a decision threshold. The GLAM is rooted in the class of linear stochastic race models (Tillman, 2017; Usher, Olami, & McClelland, 2002). These models naturally generalize to choice scenarios with more than two items and remain analytically tractable, allowing for more complex applications (e.g., embedding in a hierarchical Bayesian framework).

**Figure 3:**
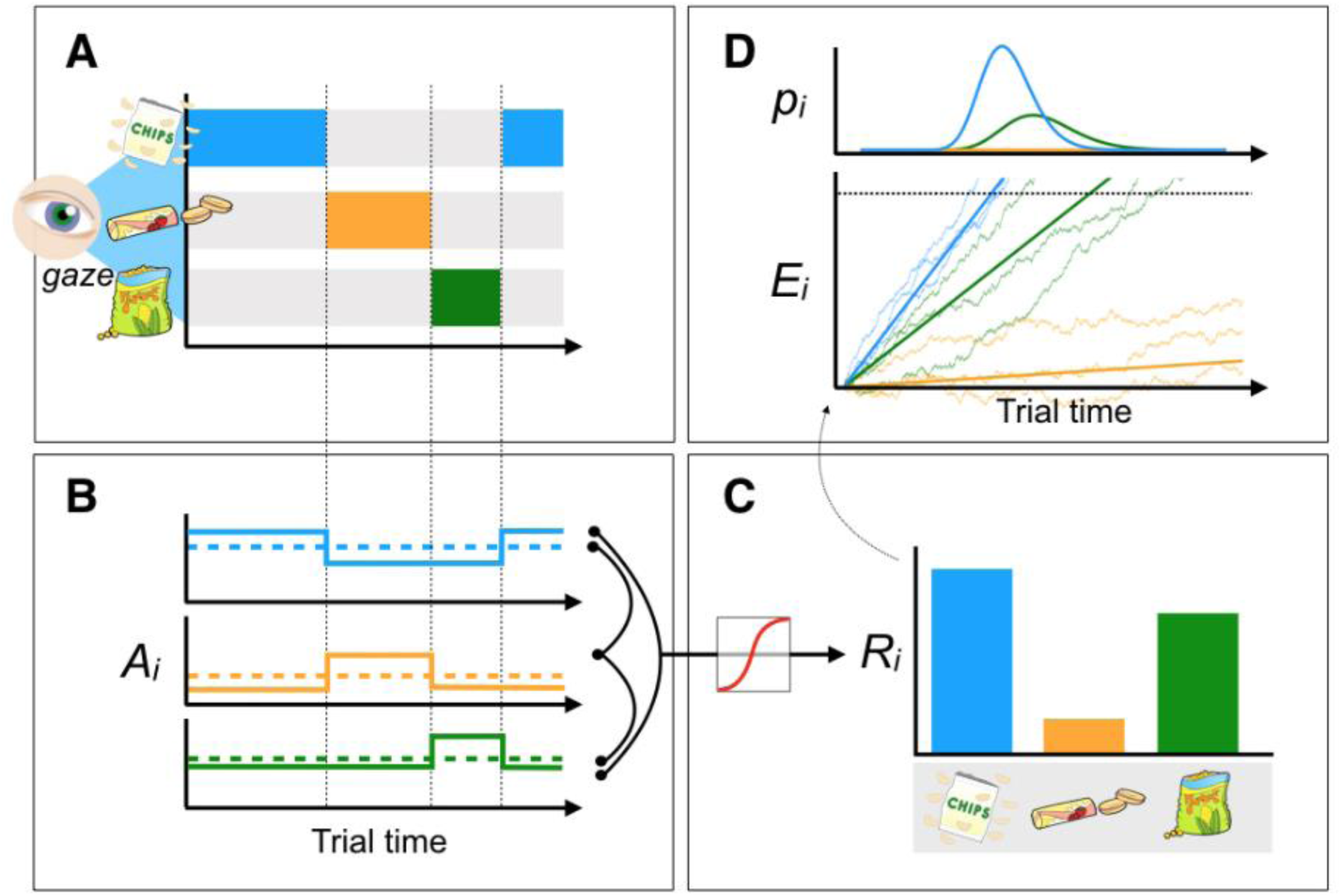
Gaze-weighted linear accumulator model (GLAM). The GLAM describes the influence of gaze allocation on the decision process, in the form of a linear stochastic race: It assumes that individuals accumulate evidence in favor of each item *i* and make a choice as soon as the relative evidence *E_i_* in favor of one item reaches a choice threshold (**D**). Importantly, the speed of the accumulation process is dependent on the distribution of visual gaze during the decision (**A & B**). For each option in the choice set, an absolute evidence signal *A_i_* is computed. The magnitude of this signal is dependent on the allocation of visual gaze, with lower magnitudes for options that are momentarily not fixated on. Absolute evidence signals are transformed into relative decision signals *R_i_* (indicating relative item preferences) by (i) computing the average absolute evidence signal for each item in the trial (dashed lines in **B**), (ii) and then computing the difference between each of these averages and the maximum of the respective other two. (iii) The GLAM assumes an adaptive representation of these relative evidence signals that is maximally sensitive to small differences in the relative decision signals. To this end, a sigmoid transform is applied (**C**). The resulting scaled relative evidence signals determine the drift terms *R_i_* in the stochastic race (**D**). The stochastic race provides first-passage time distributions *p_i_*, describing the likelihood of each item being chosen at each time point. See Methods for a more detailed model description.

In addition to the gaze bias parameter γ, the GLAM includes a general velocity parameter *v*, a noise parameter σ and a scaling parameter τ (see Methods for full model implementation details).

### Individual model comparison

We fitted and compared two GLAM variants to the response time and choice data of each participant to gauge the evidence in favor of the previously described gaze bias mechanism and to quantify its strength on an individual level:

1. A *full* GLAM variant with free parameters *v*, γ, σ, τ. This model allowed the gaze bias parameter γ to vary freely between the individuals.
2. A *no-gaze-bias* GLAM variant, where the gaze bias parameter γ was fixed to 1 (resulting in no influence of gaze on the accumulation process)

The two models differ in their complexity: The full model has one more free parameter and can therefore be expected to provide a better absolute fit to the data. We used the Deviance Information Criterion (DIC; Spiegelhalter, Best, Carlin, & Van Der Linde, 2002; Spiegelhalter, Best, Carlin & Van Der Linde, 2014) to perform model comparisons at the individual level as it includes a penalty for model complexity: more complex models are only preferred only if their added complexity is justified by an improvement in absolute fit. Lower DIC scores indicate a better model fit accounting for differences in model complexity.

The full model fitted 26 of 30 (87%) participants better than the no-gaze-bias model (Figure 4A, B). The mean ± s.e.m. difference in the DIC scores between the full and no-gaze-bias models was −34.22 ± 6.84 (Figure 4B).

**Figure 4:**
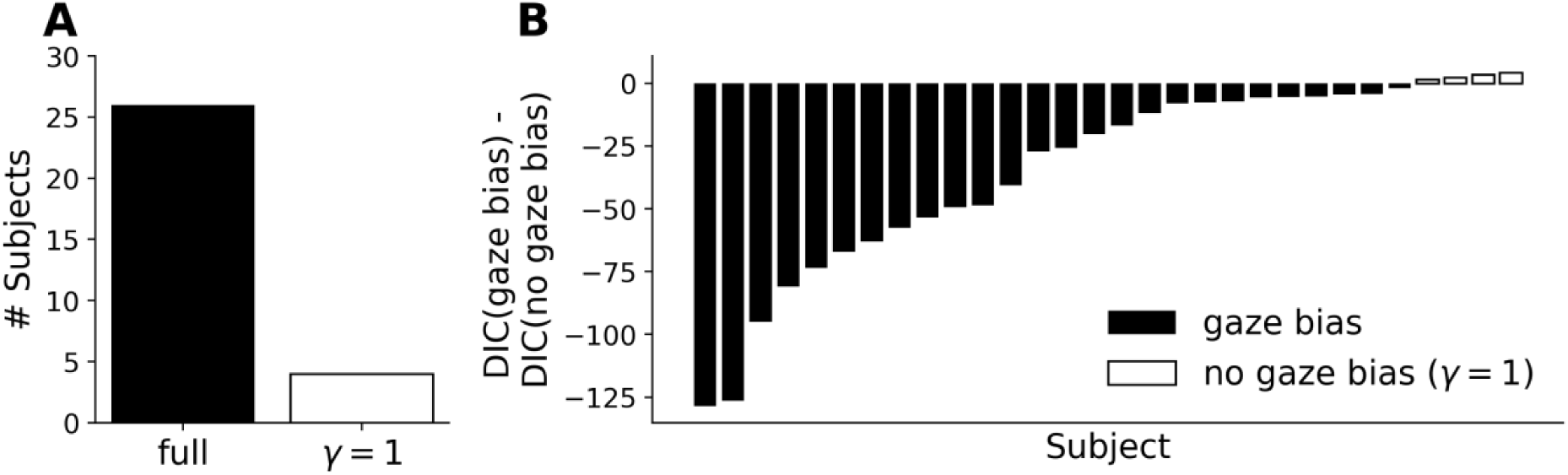
Model comparison between the full GLAM model and a no-gaze-bias GLAM (γ = 1) variant. **A**: Individual best fitting models, given by the lowest DIC score. Twenty-six of 30 participants (87%) were better described by the full model that includes a gaze bias mechanism. **B**: Individual differences in the Deviance Information Criteria (DIC) between the full and (γ = 1) model. Negative differences in the model DIC scores indicate better fits of the full model.

Individual estimates of the gaze bias parameter γ in the full model ranged from −0.93 to 0.81, with a mean ± s.d. = 0.20 ± 0.39 (Figure S2). Notably, the individual estimates covered a wide range of possible values between γ = −1 (strong gaze bias) to γ = 1 (no gaze bias). With a strong gaze bias, the GLAM leaks evidence for an item, while another is fixated on, whereas evidence accumulation is independent of gaze allocation when no gaze bias is present.

Taken together, the individual model comparison revealed that most participants’ behavior was better described by a model that includes the gaze bias mechanism. Importantly, the extent to which the accumulation process was influenced by gaze, as captured by individual gaze bias (γ) estimates, showed non-trivial individual differences.

### GLAM predicts individual choice behavior

We found that in a relative model comparison the full GLAM best describes the data of most participants, when compared to a restricted variant with no gaze bias (γ = 1; see Figure 4). However, this analysis did not take into account whether the GLAM also accurately predicts individuals’ behavior on an absolute level. To test this, we again used both model variants to simulate response data for each individual participant. This time, however, we split the data into even- and odd-numbered trials. We then used all the even trials to estimate the model parameters (training). Subsequently, we predicted the choices and response times for all the odd-numbered trials (test). The purpose of this out of sample prediction was to validate the individually estimated parameters, by comparing the GLAM’s predictions to response data that did not inform the parameter estimates. To compensate for the resulting loss of training data we fitted both GLAM variants using a hierarchical Bayesian framework (Kruschke, 2014; Wiecki, Sofer, & Frank, 2013). Here, individual model parameters and their group distributions are simultaneously estimated from the data. This is desirable as individual parameter estimates are informed by their distribution at the group level, thereby capitalizing on the information that is shared across individuals. With the hierarchical parameter estimates we then tested whether individual behavioral patterns across the three metrics are accurately predicted by the two models (see Figures 2 & 5).

**Figure 5:**
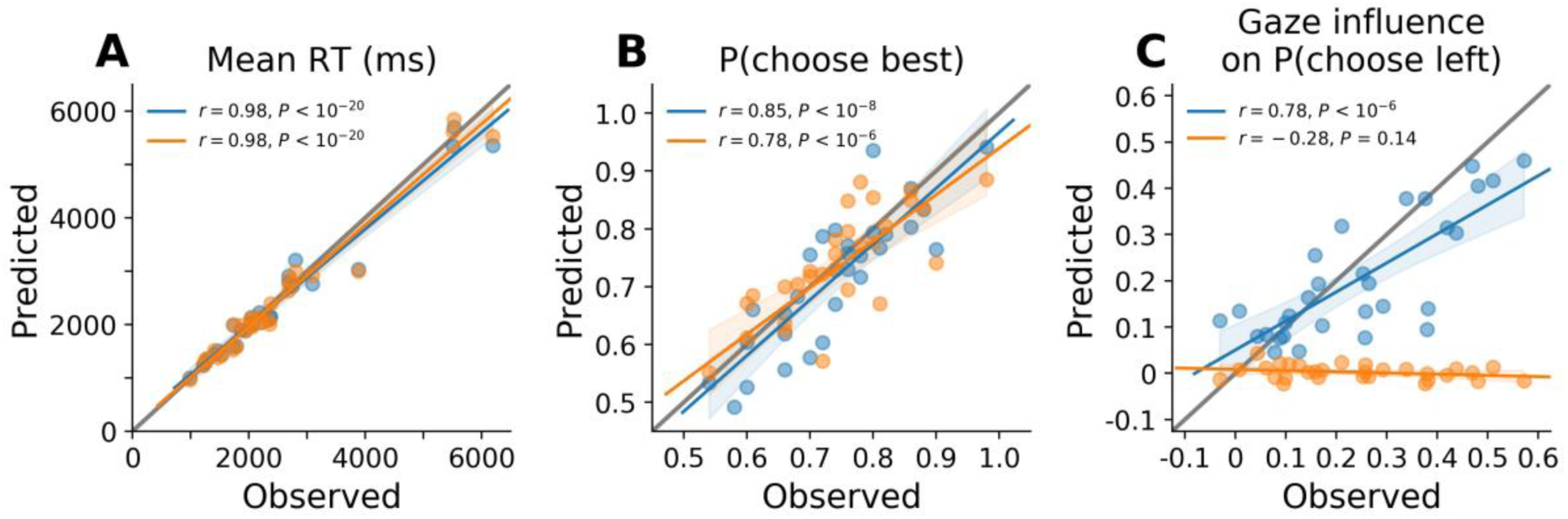
Out of sample correlations between the observed and predicted individual behavior in the odd experiment trials (the predictions were based on parameters estimated from even experiment trials). **A**: mean response time. **B**: probability of choosing the highest rated item. C: influence of gaze on choice probability. Model predictions are simulated from hierarchically estimated parameter estimates. Blue color indicates predictions from the full GLAM, whereas orange indicates predictions from a restricted GLAM variant with no gaze bias (γ = 1).

We found that the full GLAM variant accurately predicted individual differences in response times (β = 1.06, *t*(28) = 26.17, *P* < 10^−20^; Figure 5A). Similarly, individual differences in the probability of choosing the highest rated item were predicted precisely (β = 0.75, *t*(28) = 8.57, *P* < 10^−8^; Figure 5B). Lastly, we also found that the full GLAM predicted individual differences in the influence of gaze on choice probability well (β = 0.98, *t*(28) = 6.69, *P* < 10^−6^; Figure 5C).

The restricted GLAM variant with no gaze bias predicted the participants’ individual response times and the probability of choosing the highest rated item similarly well (RT : β = 1.02, *t*(28) = 27.07, *P* < 10^−20^, Figure 5A; P(choose best): β = 0.75, *t*(28) = 6.53, *P* < 10^−6^, Figure 5B). However, the model failed to predict the influence of gaze on the participants’ choices (β = −2.97, *t*(28) = −1.53, *P* = 0.14; Figure 5C), resulting in no correlation between the predicted and empirical data in our gaze influence measure.

These results showed that the full GLAM outperformed the restricted model variant in accurately predicting the participants’ empirical choices, as it also captured empirical choice patterns that are driven by gaze and not solely by the items’ liking rating.

### GLAM explains individual choice behavior

We found that the GLAM accurately predicted individuals’ response behavior. Next, we tested whether the individual model parameters are able to explain variability in the participants’ choice behavior. Here, we used standard OLS regressions to predict the three behavioral metrics in the odd-numbered trials from the individual GLAM parameters that were previously estimated hierarchically from the even-numbered trials (see Figure 6). We found that *v* (velocity parameter; see Methods for details) scaled logarithmically with the participants mean response time (β = −0.89, *t*(28) = −18.87, *P* < 10^−16^; Figure 6A). We did not find a meaningful relationship between the individual σ estimates and the probability of choosing the highest rated item (β = −1.59, t(28) = −0.19, *P* = 0.85), even though the σ parameter determines the magnitude of noise in the accumulation process (see Methods for details). However, we found that γ (gaze bias) estimates predicted the participants’ probabilities of choosing the highest rated item (β = 0.18, *t*(28) = 4.98, *P* < 10^−4^; Figure 6B), so that stronger gaze biases (smaller γ) were associated with more choices that were inconsistent with the item ratings. This relationship can be explained as follows: the gaze bias parameter γ allows the model to bias the choice process according to the distribution of gaze between items: with a strong gaze bias, the model’s predictions are strongly dependent on the distribution of gaze, and a gaze distribution that is random with respect to the items’ liking ratings then leads to random choices. On the other hand, the model’s predictions are independent of gaze when no gaze bias is present. The model then neglects gaze and predicts choices solely driven by liking ratings. Lastly, as expected, we also found that γ estimates predicted the participants’ individual scores of gaze influence on choice probability (β = −0.31, *t*(28) = −5.68, *P* < 10^−5^; Figure 6C).

**Figure 6:**
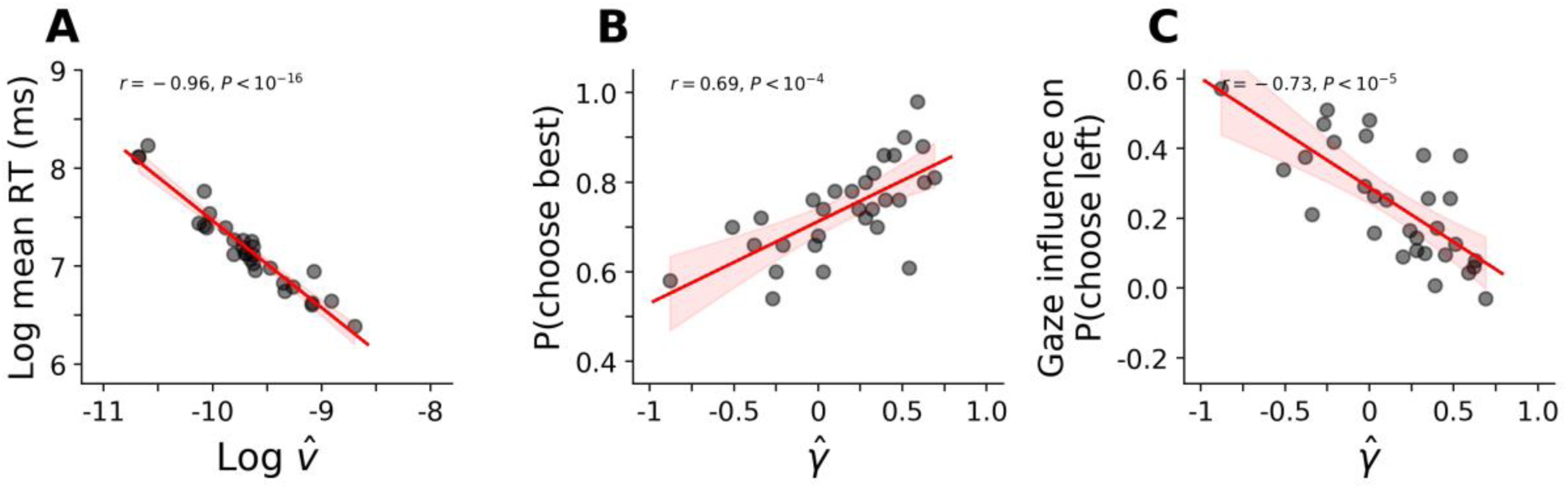
Correlations between individuals response behavior in the odd-numbered trials and the model parameters estimated from the even-numbered trials. **A**: Response time and *v* (plotted on a log-log-scale). **B**: Probability of choosing the highest rated item and γ. **C**: Influence of gaze on choice probability (mean increase in choice probability for an item that is fixated on longer than the others, when corrected for the influence of the item’s relative value and the range of the other items’ values) and γ.

## Discussion

Here, we investigated individual differences in the influence of gaze allocation on simple economic choice behavior by analyzing a previously published data set, where individuals made choices between three snack food items. We found that individuals showed an overall positive relationship between gaze and choice (longer gaze increases choice probability), but that the strength of this relationship was highly variable across individuals. To better understand the computational mechanism underlying this effect and its variability, we proposed a new model called the Gaze-weighted Linear Accumulator Model (GLAM). It assumes that individuals accumulate evidence in favor of each available item and make a choice as soon as the cumulative evidence for one item reaches a choice threshold. Importantly, the accumulation process is biased by gaze behavior, with discounted accumulation rates for unattended items. We found that the GLAM accurately predicts individuals’ choice and response time data and does so better than a model that does not assume any influence of gaze allocation on choice. We also found that the GLAM’s gaze bias estimates reliably explained individual differences in choice behavior, namely, the strength of the individuals’ association of gaze and choice behavior and the individuals’ probability of choosing the highest rated item in a choice set (stronger gaze biases were generally associated with more choices that were inconsistent with item ratings).

With the GLAM, we have provided a model that captures individual choice behavior in simple economic choice tasks with multiple alternatives with high predictive accuracy by integrating information about the individuals’ allocation of gaze. It is statistically and computationally tractable, making it readily extendable to novel choice tasks and research questions.

Our individual model comparison revealed the added value of a gaze bias mechanism in decision models. The large majority of the participants were better described by the full model, compared to a restricted variant without any influence of gaze on choice. One reason for this superior performance is that the GLAM’s use of the individual trial gaze data allows the model to make different predictions across otherwise identical choice sets. In this way, the model is able to explain variance in behavior that would otherwise be attributed to unspecified decision noise. As decision making can be seen as a stochastic process (Rieskamp, 2008), choices across identical trials with high difficulty (where ratings for the available items are very similar, for example, 3, 2 and 2 for the left, middle and right item, respectively) can be assumed to vary. A stochastic choice model without a gaze bias mechanism will make probabilistic, but identical predictions for two such trials (Gluth & Rieskamp, 2017). Leveraging a gaze bias mechanism, however, allows a model to make trial specific predictions, and these, relying on the generally positive relationship between gaze and choice, will have higher accuracy. We expect the gaze bias mechanism to be especially relevant in high-difficulty trials, where item rating information by itself provides little evidence in favor of any one alternative.

Naturally, the inclusion of gaze information into choice models leads to the question of what drives gaze during choice. This question has received considerable attention in previous research, confirming influences of item surface size (Lohse, 1997; Wedel & Pieters, 2008), position (Chandon, Hutchinson, Bradlow, & Young, 2009) and saliency (Itti & Koch, 2000) on gaze. Still, a formal integration with computationally formalized choice models was achieved only recently (Towal et al., 2013). We believe that the development of generative models for the fixation process itself, and their integration with choice models, has the potential to largely improve existing models.

Our analyses also confirmed the need for individual model fits: we found substantial variability across individuals in the influence of gaze on choice that was hidden in the group level analyses. Given that the influence of gaze on choice is variable between participants, a single gaze bias parameter γ for the whole group would not fit all the individuals well and would therefore result in inferior predictive performance of the model: participants whose link between gaze allocation and choice behavior is weaker than the group average would falsely be predicted to make choices less consistent with their value ratings, and driven more by looking behavior. Predictions for the participants’ choices with a stronger link than the group average, on the other hand, would not contain enough influence of gaze. Accounting for individual differences in the link between gaze allocation and choice behavior opens important avenues for future research, focusing on the specific determinants of these differences. For example, are these differences best characterized as a trait (stable within a person, but variable between persons), state (variable within a person, between different situations or contexts) or both (variable between persons and contexts) (see Peters & Büchel, 2011, for a similar discussion in the context of delay discounting)?

Despite a wealth of findings exploring the computational mechanisms underlying simple choice behavior and its link to visual fixations (e.g., Shimojo et al., 2003; Armel et al. 2008; Krajbich et al., 2010; Krajbich & Rangel, 2011; Towal et al., 2013; Cavanagh et al., 2014; Fisher, 2017), most of this work, and the associated computational frameworks (e.g., Ratcliff et al., 2016), is difficult to extend to complex choice scenarios (i.e., involving more than two choice alternatives). Here, we have shown that the GLAM captures individuals’ choice behavior well in choice situations with only few choice alternatives. However, the GLAM naturally extends to choices involving many more options, as we mostly encounter in our everyday lives. Imagine standing in front of a vending machine to buy a snack. These machines can easily store up to 20 items. We assume that in these multialternative choice situations, both gaze and individual differences will play an even more prominent role: individuals, when confronted with large choice sets, do not always look at all available items (e.g., Reutskaja, Nagel, Camerer, & Rangel, 2011). A choice model that considers individuals’ liking values only will therefore fail in accurately predicting individuals’ choice behavior. A model that includes information about individuals’ gaze distribution during decision formation will, on the other hand, outperform such naive models, because it will better account for the set of items that individuals actually consider for a choice. In addition, we assume that behavioral differences between individuals to increase with increasing choice set size. For example, we assume that some individuals may look at only a few of the available items, before making a choice, while others may spend a long time searching for the most highly valued option (as indicated in Reutskaja et al., 2011). To understand whether there is a common choice mechanism underlying these different types of choice behavior, it is necessary to test the ability of a model to capture individual choice patterns.

Real life choices have another level of complexity often not considered in simple economic choice tasks; options comprise multiple, oftentimes orthogonal attributes. For example, each item in a vending machine is associated with a price that has to be considered. Consumer goods often also include other attributes (e.g., ratings by other consumers: De Martino, Bobadilla-Suarez, Nouguchi, Sharot, & Love, 2017; energy consumption, etc.), which are commonly displayed visually to the decision maker. Certain configurations of option attributes can induce systematic shifts in preference, called context effects (Mohr, Heekeren, & Rieskamp, 2017; Soltani, De Martino, & Camerer, 2012; Trueblood, Brown, Heathcote, & Busemeyer, 2013). These preference shifts can vary considerably between individuals (Mohr et al., 2017). Notably, eye-tracking data about the identity and sequence of fixated attributes are predictive of choice in context effect settings (Noguchi & Stewart, 2014). Future research on these effects and their relationship to gaze requires a model that can be fitted on an individual basis and is applicable to multialternative choice scenarios. The GLAM provides a starting-point to explore these types of research questions in the future.

Like other existing decision making models (e.g., Usher and McClelland, 2001; Roe, Busemeyer & Townsend, 2001; Ratcliff et al., 2016; Krajbich et al., 2010), the GLAM also incorporates several assumptions about the neural computations underlying simple economic choices. It is necessary to evaluate the plausibility of these assumptions, next to the ability of a model to capture individuals’ choice behavior. The GLAM assumes evidence accumulation towards a decision threshold and a fixation-dependent bias of this process. There is strong neural evidence for accumulation-to-bound processes during decision formation in a variety of choice tasks (e.g., Basten, Biele, Heekeren, & Fiebach, 2010; Philiastides & Sajda, 2007; Churchland, Kiani & Shadlen, 2008; Liu & Pleskac, 2011; O’Connell, Dockree & Kelly, 2012; Wyart, De Gardelle, Scholl & Summerfield, 2012; Polanía, Krajbich, Grueschow & Ruff, 2014; Lafuente, Jazayeri & Shadlen, 2015; for a review see Gold & Shadlen, 2001, and Heekeren, Marrett & Ungerleider, 2008). Recently, it was also shown that single-trial EEG components reflecting attention in simple perceptual decision-making tasks explain variance in single-trial evidence accumulation rates of the decision process (Nunez, Vandekerckhove,& Srinivasan, 2017) and that variability in these components can explain behavioral differences between individuals (Nunez, Srinivasan, & Vandekerckhove, 2015). Two recent studies also provided first empirical evidence that value-driven activity in the orbitofrontal cortex of monkeys is modulated by fixation location when they viewed reward-associated visual cues in a free-viewing paradigm (Hunt, Malalasekera, Berker, Miranda, Farmer, Behrens & Kennerley, 2017; McGinty, Rangel, & Newsome, 2016). Together, these studies provided the first neurobiological evidence of the influence of visual fixations on the process of decision formation. Ultimately, a better understanding of these computations will be central to building holistic models of the choice process and for advancing existing choice frameworks. In addition, it might also help us to better understand the origin of behavioral variability that we observe within and between individuals.

The focus on individual differences in the relationship between gaze and choice behavior can also prove itself relevant in clinical research domains. Increasingly prevalent clinical conditions, such as type 2 diabetes and obesity, typically involve maladaptive decision-making between visually presented stimuli, often strategically positioned and designed to capture attention. Snack food items, for example, are advertised with bright, salient colors and placed prominently (e.g., at eye level, near the checkout in the supermarket), which could have adverse effects on individuals prone to making maladaptive food choices. Healthier diets (i.e., food choices) are both prevention and treatment for such diseases, and a better understanding of how individuals’ decisions are impacted by looking behavior could help inform the search for predictors of clinical behavior and improve therapeutic approaches. In addition, individually tailored therapeutic approaches, based on a better understanding of individual response patterns, promise higher efficacy and, in turn, reduced health care spending.

## Methods

### Data, tasks, procedure & preprocessing

We reanalyzed a data set that was previously published in Krajbich and Rangel (2011). In the corresponding experiment, hungry participants made repeated choices between multiple snack food items (e.g., Twix, Lays Chips, Skittles, etc.), while their eye movements were recorded.

The data set contains data from 30 Caltech students, who reported to regularly eat the snack foods that were used in the experiment and had no dietary restrictions. The participants received a show-up fee of $20 and one food item. The experiment was approved by Caltech’s Human Subjects Internal Review Board.

All participants were asked not to eat for 3 hours prior to the experiment. In an initial liking rating task the participants indicated liking ratings between −10 to 10 for each of the 70 different snack food items using an on screen slider with a randomized starting point and free response time (“How much would you like to eat this at the end of the experiment?”; Figure 1, Task 1). These ratings were used as a measure of the value participants placed on each item. In the subsequent choice task the participants made choices between triplets of food items. The items were arranged in a triangular fashion on the screen (Figure 1, Task 2). In one half of the trials, this triangle pointed upwards (center option on top), in the other half it pointed downwards (center option at the bottom). Choices were indicated with free response times and using the left, down and right arrow keys on a keyboard. Each trial began with a 2 s forced fixation towards the center of the screen. A yellow feedback box was shown around the chosen item for 1 s after a choice was made. Lastly, the participants were required to stay for 30 min after the experiment, to eat a food item that they chose in one randomly selected choice trial. The participants performed 100 choice trials each.

The participants’ eye movements were continuously recorded with a 50 Hz desktop-mounted Tobii eye tracker.

The data were obtained from the original authors in an already preprocessed format. The original preprocessing steps included the removal of trials with missing fixation data for more than 500 ms at the beginning or end of the trial, resulting in a total of 2966 remaining trials (mean ± s.e.m. number of trials dropped per participant was 1.1 ± 0.9). Rectangular areas of interest (AOIs) were constructed around each food item in each trial and visual fixations were assigned to the corresponding item or coded as non-item fixations. If a non-item fixation was preceded and succeeded by fixations on the same item, the non-item fixation would also be assigned to this item. Other non-item fixations were not reassigned and discarded from all further analyses.

### Gaze-weighted Linear Accumulator Model (GLAM) details

The GLAM belongs to the class of linear stochastic race models (Usher & McClelland, 2001). It assumes accumulation of noisy evidence in favor of each available alternative *i*, and that the choice is determined by the first accumulator that reaches a common boundary. In particular, we define the accumulated relative evidence *E_i_* in favor of alternative *i*, as a stochastic process that changes at each point in time *t* according to:

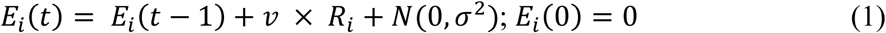

*E_i_* consists of two separate components: a drift term *R_i_* and zero-centered normally distributed noise with standard deviation σ. The overall speed of the accumulation process is governed by the velocity parameter *v*. The drift term *R_i_* describes the average amount of relative evidence for item *i* that is accumulated at each point in time *t*. We define the relative evidence 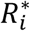 as the difference in the stationary absolute evidence signal *A_i_* of item *i* and the maximum absolute evidence of all other items *J*:

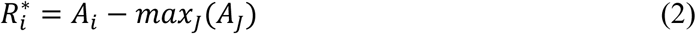

The model’s gaze bias mechanism is implemented in the absolute evidence signal *A_i_*: Similar to the aDDM, the absolute evidence signals are assumed to be proportional to the value ratings *r_i_*, and crucially, switch between two different states during the trial: an unbiased state, when an item is currently looked at, and a biased state, when gaze is directed towards a different item. Therefore, on average, *A_i_* is a linear combination of two terms that are weighted by the fraction *g_i_* of the total fixation time that item *i* was fixated in the trial:

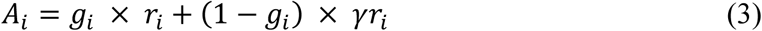

Here, γ (−1 ≤ γ ≤ 1) is the model’s gaze bias parameter that determines the strength of the downweighting during the biased state. If γ = 1, there is no difference between the biased and unbiased state, producing no gaze bias. If γ < 1, the absolute evidence signal is discounted by the γ parameter, resulting in a gaze bias. If −1 ≤ γ < 0, the sign of the evidence signal changes, thereby leaking evidence, when the item is not fixated. This leakage mechanism is supported by a recent empirical study (Ashby et al., 2016). The maximum amount of evidence that can be accumulated or leaked at each time point is symmetric in magnitude, as the γ parameter is bounded between −1 and 1.

Note that the range of possible 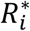 (equation (2)) depends on the participants’ use of the item rating scale: if the ratings only cover a narrow range of possible values on the scale, the relative evidence values 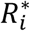 will likewise be small, whereas they will be large if the participant utilizes the entire range of the rating scale. GLAM assumes an adaptive representation of the relative evidence signals that is compensating for the participants’ use of the rating scale and thereby sensitive to marginal differences in the relative evidences, particularly to values close to 0 (where the absolute evidence signal for one item is only marginally different to the maximum of all others). To this end, a logistic transform *s(x)*, with scaling parameter τ is applied:

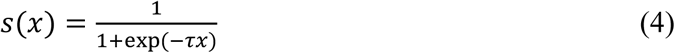

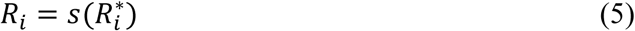

The first passage time density *f_i_(t)* of a single linear stochastic accumulator *E_i_*, with decision boundary *b*, is given by the Inverse Gaussian Distribution (Wald, 1973):

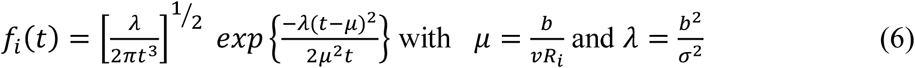

However, this density does not take into account that there are multiple accumulators in each trial racing towards the same boundary. As soon as any of these accumulators crosses the boundary a choice is made and the trial ends. For this reason, *f_i_(t)* must be corrected for the probability that any other accumulator crosses the boundary first. The probability that a single accumulator crosses the boundary prior to *t*, is given by its cumulative distribution function *F_i_(t)*:

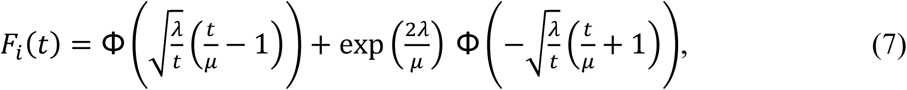

where Φ(x) is the standard normal cumulative distribution function. Hence, the joint probability *p_i_(t)* that accumulator *E_i_* crosses *b* at time *t*, and that no other accumulator *j* has reached *b* first, is given by:

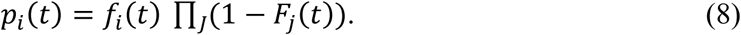

We performed a parameter recovery study to rule out misspecifications of the model and assert the validity of the parameters estimated from empirical data. All of the parameters could be recovered to a satisfying degree (see Figure S1 for detailed results).

Although the race framework deviates from the classical Drift-Diffusion Model (DDM; Ratcliff, 1978; Ratcliff et al., 2016), which is known to implement an optimal decision procedure in the sense of the sequential probability ratio test (Bogacz, Brown, Moehlis, Holmes, & Cohen, 2006), it has reasonable benefits in the context of this paper: first, the model naturally generalizes to choices between three alternatives, which is not trivial for the classical DDM. Second, it generalizes to settings with even larger choice sets. Third, an analytical solution for the first-passage time density of the linear stochastic race exists. This solution enables a very fast and efficient parameter estimation, without the need to numerically estimate densities from a large number of model simulations.

### GLAM parameter estimation

#### Individual

For the individual model comparison, we first estimated the model parameters at the individual level. The full GLAM has four parameters (*v*, γ, σ, τ). The individual models were implemented in the Python library PyMC3 (Salvatier, Wiecki, & Fonnesbeck, 2016) and fitted using the default Markov Chain Monte Carlo No-U-Turn-Sampler (NUTS; Hoffman & Gelman, 2014). We parameterized the model so that the noise parameter σ was sampled proportionally to the velocity parameter *v* using a signal-to-noise variable *SNR* (note that this does not add a free parameter to the model, as σ is now fully determined by *v* and *SNR*). We placed uninformative, uniform priors between sensible limits on all model parameters:

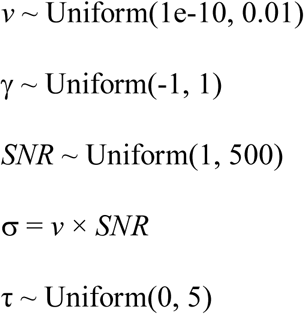

Further, we assumed a fixed 5% rate of error trials, which we model as a participant-specific uniform likelihood distribution *u_s_(t)*. This error likelihood describes the probability of a random choice for any of the *N* available choice items at a random time point in the interval of empirically observed response times (cf. Ratcliff & Tuerlinckx, 2002; Wiecki et al., 2013):

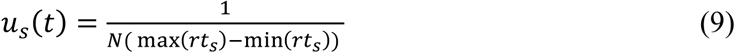

The resulting choice likelihood is then given by:

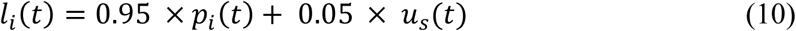

For each individual model, the NUTS sampler was initialized using the default behavior in PyMC 3.2, followed by 500 tuning samples that were discarded. Finally, we drew 2000 posterior samples that we used to estimate the model parameters.

In addition, a restricted no-gaze-bias GLAM variant was also fit to the individual data. It was specified and fitted identically to the full model, but had the gaze-bias parameter γ fixed at 1.0. The reported parameter estimates are maximum a posteriori (MAP) estimates.

#### Hierarchical

We also estimated the GLAM parameters in a hierarchical Bayesian framework (Kruschke, 2014; Vandekerckhove, Tuerlinckx, & Lee, 2008; Wiecki et al., 2013). Here, the fit of participant level parameters is informed by the group distribution of parameters. Each parameter on the participant level is modeled as coming from a population distribution whose shape and location are also estimated from the data. We assumed that all participant level parameters are drawn from normal population distributions, which we bounded to sensible ranges:

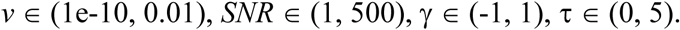

We used ADVI (Kucukelbir, Ranganath, Gelman, & Blei, 2015) from the PyMC3 python library (Salvatier et al., 2016) to approximate the posterior distribution of the model parameters. In analogy to the individual models, we assumed a fixed 5% rate of participant-specific uniformly distributed error responses (see above). The reported parameter estimates are maximum a posteriori (MAP) estimates.

### Model simulations

Choice and response time data was simulated from the GLAM according to the following procedures: each trial in the left-out data set, containing all the odd-numbered trials, was repeated 50 times. For every trial the model used the empirically observed item ratings and gaze distributions. With a fixed rate of 5% the simulation produced a random choice and a random response time between the participant’s minimum and maximum observed response times (cf. equations (9) & (10)). With a rate of 95% the choice and response time were simulated from the actual GLAM: for each item in the trial, a first passage time (FPT) was drawn according to the single-item first passage densities (equation (8)). The response time and choice were then determined by the item with the shortest FPT.

### Availability of data, model and analysis code

All analyses and figures can be reproduced using the data set, scripts and GLAM resources that are available at http://www.github.com/glamlab/glam.

#### Software

All analyses were performed in Python, using the NumPy and SciPy (Van der Walt, Colbert & Varoquaux, 2011), Pandas (McKinney, 2010), Statsmodels (Skipper & Perktold, 2010), PyMC3 (Salvatier et al., 2016) and Theano (Theano Development Team, 2016) libraries. We used Matplotlib (Hunter, 2007) for visualization.

## Acknowledgements

Peter N. C. Mohr is funded by the Freie Universität Berlin within the Excellence Initiative of the German Research Foundation (DFG). Further funding is provided by the WZB Berlin Social Science Center. Armin W. Thomas is funded by the Bernstein Center for Computational Neuroscience Berlin (BCCN). Felix Molter is supported by the International Max Planck Research School on the Life Course (LIFE). Ian Krajbich is funded by National Science Foundation Career Award 1554837.

## Author contributions

A.W.T, F.M., H.R.H. and P.N.C.M. conceived the study. A.W.T. and F.M. designed the model and carried out the data analysis. A.W.T, F.M., I.K., H.R.H. and P.N.C.M. cowrote the manuscript.

## Supplementary Information

### Parameter recovery

**Figure S1:**
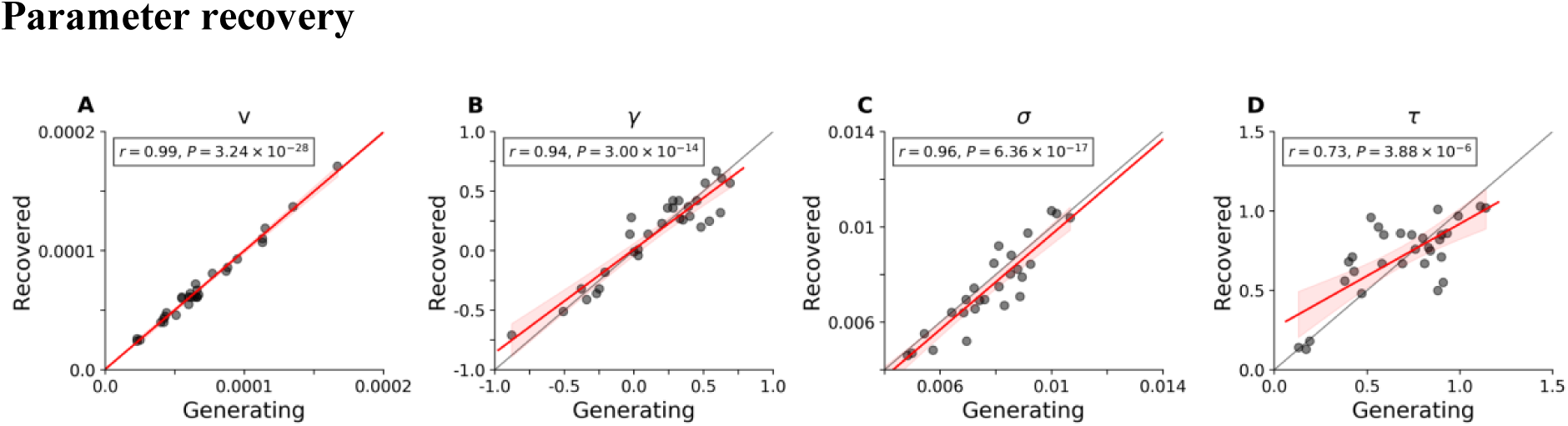
Results of a parameter recovery study of the GLAM. Parameter estimates were estimated from model simulated data sets. Panels **A** to **D** show relationships between generating and recovered parameters. All parameters could be recovered to a satisfying degree.

We performed a parameter recovery study to validate the parameter estimates. We simulated data using the GLAM and the corresponding hierarchically estimated individual parameter estimates. Each empirically observed even-numbered trial (i.e., set of item value ratings and relative gazes in training set trials) was simulated once, resulting in a GLAM-generated data set that matches the original training data in size and structure. We then performed the exact same hierarchical parameter estimation procedure as we did in the training data. Ideally, the recovered parameters should match the originally estimated ones. The correlations between generating and recovered parameters are displayed in Figure S1. The critical model parameters v, γ and σ were recovered very well (Figure S1 A, B, C, respectively). We found a significant linear relationship between generating and recovered τ parameters (Figure S1D), although they include some level of variance.

### Parameter estimates

**Figure S2:**
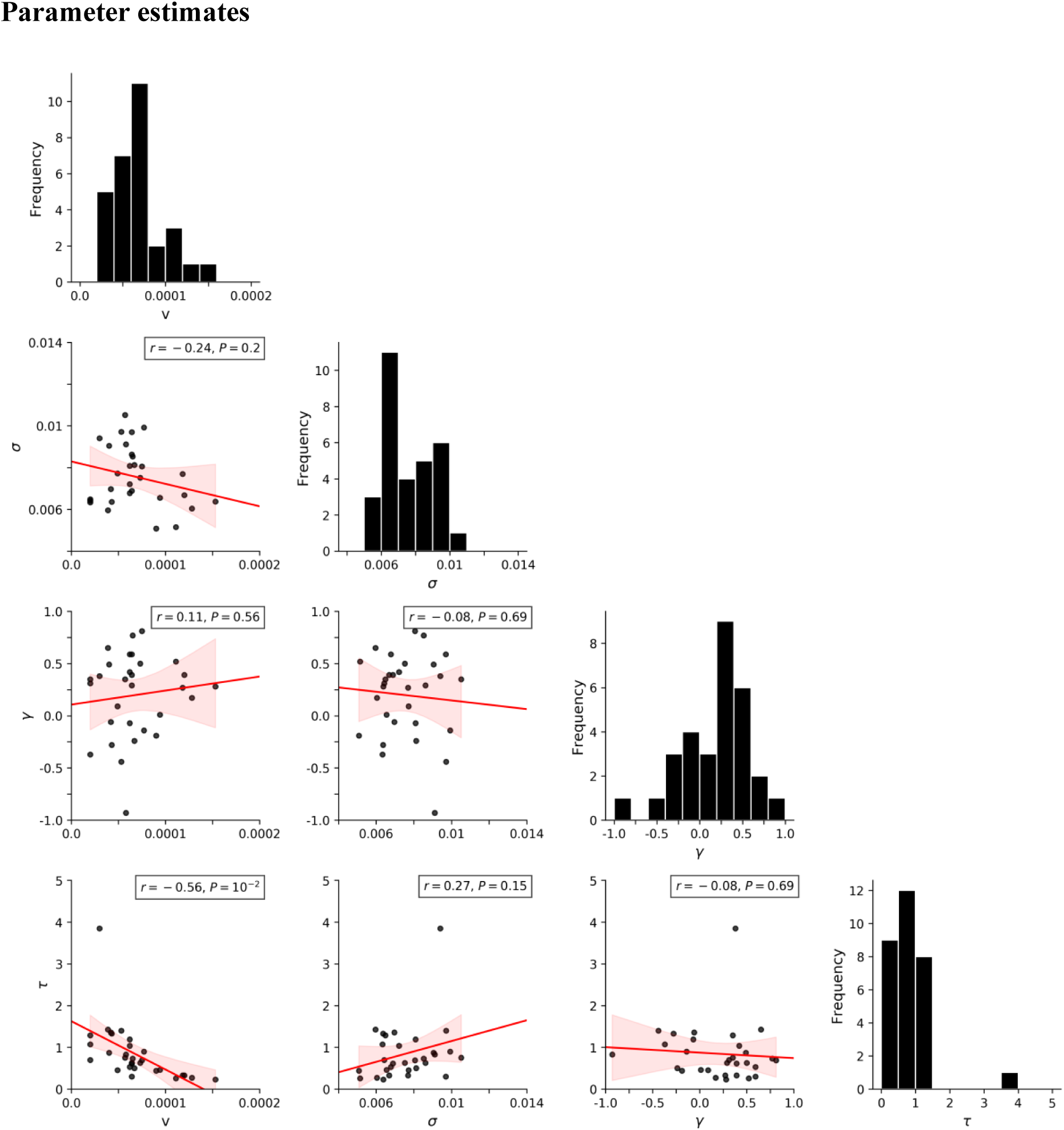
Parameter estimates and correlations between parameters. Estimates shown are from individual model fits.

